# Genetic access to functionally distinct ganglion cell types in the Somatostatin-IRES-Cre mouse retina

**DOI:** 10.1101/819441

**Authors:** James W. Fransen, Bart G. Borghuis

## Abstract

Retinal ganglion cells (GCs) are a functionally diverse neuron population that encodes and transmits distinct representations of the visual image on the retina to target nuclei in the brain. Independent studies of visually-evoked responses, cell morphology, and gene expression each suggest that GCs in mouse may comprise as many as forty distinct cell types. To date, only a subset of these types have been characterized in detail, and for most genetic access is still lacking. Thus, the majority of identified GC types remains inaccessible for targeted electrophysiology and functional imaging, precluding efficient studies of their response properties, and the cell-intrinsic mechanisms and presynaptic circuits that generate these properties. Here we show that an existing mouse line that is commonly used for studies of cortical inhibitory circuits – Somatostatin-IRES-Cre (Sst-Cre), consistently labels an understudied subset of four GC types with distinct visual responses. We characterized these types both anatomically and functionally using Cre-dependent reporter mouse lines and confocal fluorescence imaging, calcium imaging, and whole-cell electrophysiology. We show that one of the labeled GC types is suppressed by luminance contrast, while another matches a recently described orientation-selective GC type. Our results give new information about these two identified GC types, and establish the utility of the Sst-Cre transgenic mouse line for studies of recently identified GC circuits in the mouse retina.

## Introduction

The emergence of transgenic mouse lines with selective Cre-recombinase expression (Cre-driver mice) and loxP-flanked fluorescent protein gene inserts (Cre-dependent reporter mice) has provided a powerful toolset for probing neural circuits and circuit-level computations in the retina. For example, starburst amacrine cells, VIP amacrine cells, and ON-OFF direction selective (DS) GCs are experimentally accessible thanks to Cre-driver mice and much of our current knowledge about these types relied on efficient genetic access for targeted recording (Huberman et al., 2009; Yonehara et al., 2011; Park et al., 2014; Park et al., 2015; Fransen and Borghuis, 2017). However, genetic access to the majority of the retina’s ~200 neuron types is still lacking. As a consequence, ongoing efforts are aimed at identifying mouse lines with selective expression in one or small number of neuron types, which are most useful for targeted studies. Here, we report on one such line, Somatostatin-IRES-Cre (Taniguchi et al., 2011).

To represent the visual image transduced by the photoreceptors, neuronal circuits filter different aspects of the visual input into multiple parallel signaling pathways. At the level of the retinal output, the ganglion cell layer, these parallel pathways culminate in a set of spatially overlapping GC types with diverse visual responses. The most recent classification of these types in mouse distinguished 39 GC types, based on exhaustive population calcium imaging and hierarchical clustering of their visually-evoked responses and (Baden et al., 2016). A major current challenge is to explain this functional diversity at the cellular and circuit level.

Two major experimental approaches toward this goal—functional imaging and whole-cell electrophysiology—benefit directly and strongly from selective genetic access to each of the identified GC types. A recent tally lists eighteen identified GC types in existing transgenic mouse lines (Sumbul et al., 2014; Sanes and Masland, 2015). Thus, we currently still lack genetic access to about half of the defined GC types, and the search for useful GC expression patterns in existing mutant mouse lines is necessary, and ongoing.

The Somatostatin-IRES-Cre (Sst-Cre) line has been used extensively to study inhibitory interneurons in cortical neural circuits (Scheyltjens and Arckens, 2016). Inhibitory interneurons including GABAergic amacrine cells (ACs) in the retina, too, are known to express somatostatin (SST) (Larsen et al., 1990; White et al., 1990). Interestingly, crossed with a Cre-dependent fluorescent reporter line, retinae of Sst-Cre mice show fluorescent label also in a subset of (excitatory) GCs, likely through low-level *Sst* gene expression (Martersteck et al., 2017). While SST expression in mammalian retinal GCs has been reported previously (Rickman et al., 1996), whether the Cre-expressing GCs observed in Sst-Cre mice comprise a fixed and limited subset of GC types, and if so, which types they are, is not known.

To address this, we crossed the Sst-Cre mouse with Cre-dependent reporter mouse lines (Ai9, Thy1-stop-YFP, or Ai95; Buffelli et al., 2003; Madisen et al., 2010; Madisen et al., 2015) and determined the morphological and electrophysiological characteristics of genetically labeled GCs using targeted dye-fills of isolated cells, electrophysiological whole-cell recording, and two-photon fluorescence calcium imaging. While Sst-Cre labels multiple GCs types, we consistently encountered four subtypes that each accounted for between 7% and 16% of the total labeled GC population. Of these four types, two were of particular current interest. The first type functionally and anatomically matches the recently reported suppressed-by-contrast cell (SBC; (Tien et al., 2015), an apparent analog of the uniformity detector (UD) GC reported in rabbit (Sivyer et al., 2010). The second type matches one of the recently described orientation selective, non-direction selective GCs (OS GC; (Nath and Schwartz, 2017) discovered through ‘blind’ (non-targeted) recording of unlabeled GC populations. Our results demonstrate that the Sst-Cre transgenic mouse line is useful for efficient studies of a select subpopulation of excitatory neuron types in the mouse retina.

## Methods

### Animals

All animal procedures were approved by the Institutional Animal Care and Use Committee at the University of Louisville and were in compliance with National Institutes of Health guidelines. Somatostatin-IRES-Cre mice (Sst-Cre; Jackson Laboratory stock Sst^tm2.1(cre)Zjh^/J; see Taniguchi et. al, 2010) were bred with reporter mice (Ai9 (B6.Cg-*Gt(ROSA)26Sor^tm9(CAG-tdTomato)Hze^*/J; JAX 7909) or Thy1-Stop-EYFP (B6.Cg-Tg(Thy1-EYFP)15Jrs/J); JAX 5630) to fluorescently label Sst-Cre positive neurons in the retina. For calcium imaging experiments, Sst-Cre mice were bred with Ai95 mice (B6J.Cg-Gt(ROSA)26Sor^tm95.1(CAG-GCaMP6f)Hze^/MwarJ; JAX 28865) to express GCaMP6f in Sst-Cre positive neurons. Mice of either sex aged between P30 and P120 were used for experiments.

### Retinal Preparation

Mice were dark adapted for 30 minutes prior to euthanasia and tissue harvest of the retina. Animals were anesthetized using Isofluorane and euthanized via cervical dislocation. Each eye was enucleated and placed in oxygenated (95% O_2_/5% CO_2_) Ames medium (Sigma-Aldrich). The retina was dissected from the eyecup, radially incised to identify the ventral and dorsal halves, and placed on a nitrocellulose membrane filter disc (EMD Millipore) with the GC side up. Four ~1.1mm diameter holes in the filter disc, one under each quadrant of the retina permitted visual stimulation of the retina through the microscope condenser light path. The filter disc with retinal preparation was placed in a custom-designed, 3D-printed recording chamber and held in place using a clamping ring with parallel nylon wires (0.076 mm diameter; Frog Hair, 8X,; Amazon). Retinas were continuously perfused at ~4-6 mL/min with oxygenated Ames medium warmed to~34°C for the duration of an experiment.

### Cell Morphology

For morphological reconstruction, fluorescently labeled Sst-Cre positive neurons were targeted for sharp electrode fills with neurobiotin (Vector Laboratories, California). Following neurobiotin fills, the retina was fixed for 20 minutes using a 5% paraformaldehyde solution and then washed and stored in 0.1M phosphate buffered saline (PBS). The fixed retina was then reacted with Streptavidin conjugated to Alexa-488 (Thermo-Fisher Scientific) at room temperature, overnight, under continuous slow motion on an orbital shaker. Following three washes with PBS, the retina was mounted in Vectashield on a glass slide with a 0.12 mm spacer to prevent flattening (Secure Seal, SS1×9; Grace BioLabs), covered with a coverslip, and sealed with clear nail protector (Wet N Wild; Amazon). Retinas were imaged using an FV-1000 laser-scanning microscope (Olympus Life Science). Non-biotin filled fluorescent Sst-Cre neurons in the inner nuclear layer (INL) and amacrine cell layer (ACL) were used to define the outer and inner border of the IPL to compute stratification depth.

### Electrophysiology

Fluorescently labeled neurons were targeted for whole-cell recording using a custom built two-photon microscope set for emission at 925 nm. The two-photon microscope was controlled by ScanImage software, v3.8 (www.scanimage.org; (Pologruto et al., 2003). Glass electrodes (4-7.5 MΩ resistance) were pulled using a P-97 pipette puller (Sutter Instruments, California). The intracellular solution contained (in mM): 120 Cs-methanesulfonate, 5 TEA-Cl, 10 HEPES, 10 BAPTA, 3 NaCl, 2 QX-314-Cl, 4 ATP-Mg, 0.4 GTP-Na2, and 10 phosphocreatine-Tris2 (pH 7.3, 280 mOsm). Excitatory and inhibitory currents were recorded at the reversal potential for chloride (−68 mV) or cations (+15 mV), respectively, after correcting for the liquid junction potential (−9 mV). Alexa Fluor 488 fluorescent dye (Thermo Fisher Scientific) was added to the intracellular solution to fill the recorded cell during recording to obtain post-hoc dendritic arbor morphology and stratification depth using 2-photon fluorescence imaging. Membrane currents or membrane voltage were amplified using a MultiClamp 700B amplifier and digitized using a DigiData 1550, controlled with pCLAMP 11 software (all Molecular Devices, CA).

### Calcium Imaging

Fluorescence responses of GCaMP6f-expressing Sst-Cre neurons were obtained using a custom-built two-photon fluorescence microscope powered by an ultrafast pulsed laser tuned to 910 nm (Coherent Ultra II; Coherent, Santa Clara, CA.

### Light Stimulation

We used a DLP projector (Notebook Projection Companion; Hewlet-Packard, California), fitted with a UV emitting LED (λ_max_ = 395 nm) to optimally stimulate UV-opsin expressing cones in the ventral mouse retina. The stimulus was focused onto the photoreceptor layer of the retina using the microscope condenser, and image focus was verified prior to each experiment. Stimuli were generated using custom-written software and included contrast reversing spots, 1 Hz drifting square-wave gratings at eight orientations (0, 45, 90, 135, 180, 225, 270, and 315 degrees), and contrast reversing static square-wave gratings presented at 100% Michelson contrast, also modulated at 1 Hz.

### Data Analysis

Data were analyzed in Matlab (The Mathworks; Natick, MA) using custom-written algorithms. Statistical significance (p < 0.05) was assessed using Student’s T-Test. Error bars represent +/− standard error of the mean (SEM) throughout. Direction selective index (DSI) values were calculated as follows:

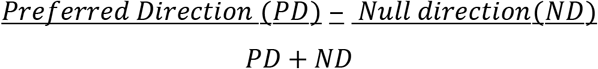

Orientation selective index (OSi) values were calculated as follows:

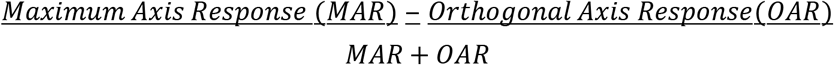

## Results

### Distribution density and morphology of Sst-Cre positive GCs

Crossing Sst-Cre transgenic mice with the Ai9 Cre-dependent fluorescence reporter mouse (Madisen et al., 2010) gave a retinal expression pattern with labeled neurons in both the GC and AC layer. Fluorescence-labeled cells in the GC layer–our sole focus here–were distributed across the entire retina with no obvious bias along the dorsal-ventral or temporal-nasal axis (Figure 1A). The number of labeled cells in animals homozygous for Sst-Cre was about 1.5-fold larger compared with animals heterozygous for Sst-Cre (3778 ± 146, n = 3 vs. 2634 ± 235, n=3; t-test, *p*=0.014), indicating that gene recombination-dependent fluorescence expression was limited by the cellular level of Cre-recombinase expression. Based on the reported total number of GCs in the mouse retina, ~60,000 (Williams et al., 1996), labeled GC populations in Sst-Cre^+/+^ and Sst-Cre^+/−^ represented approximately 6% and 4% of all GCs, respectively, which we consider a sparse label.

**Fig 1.**
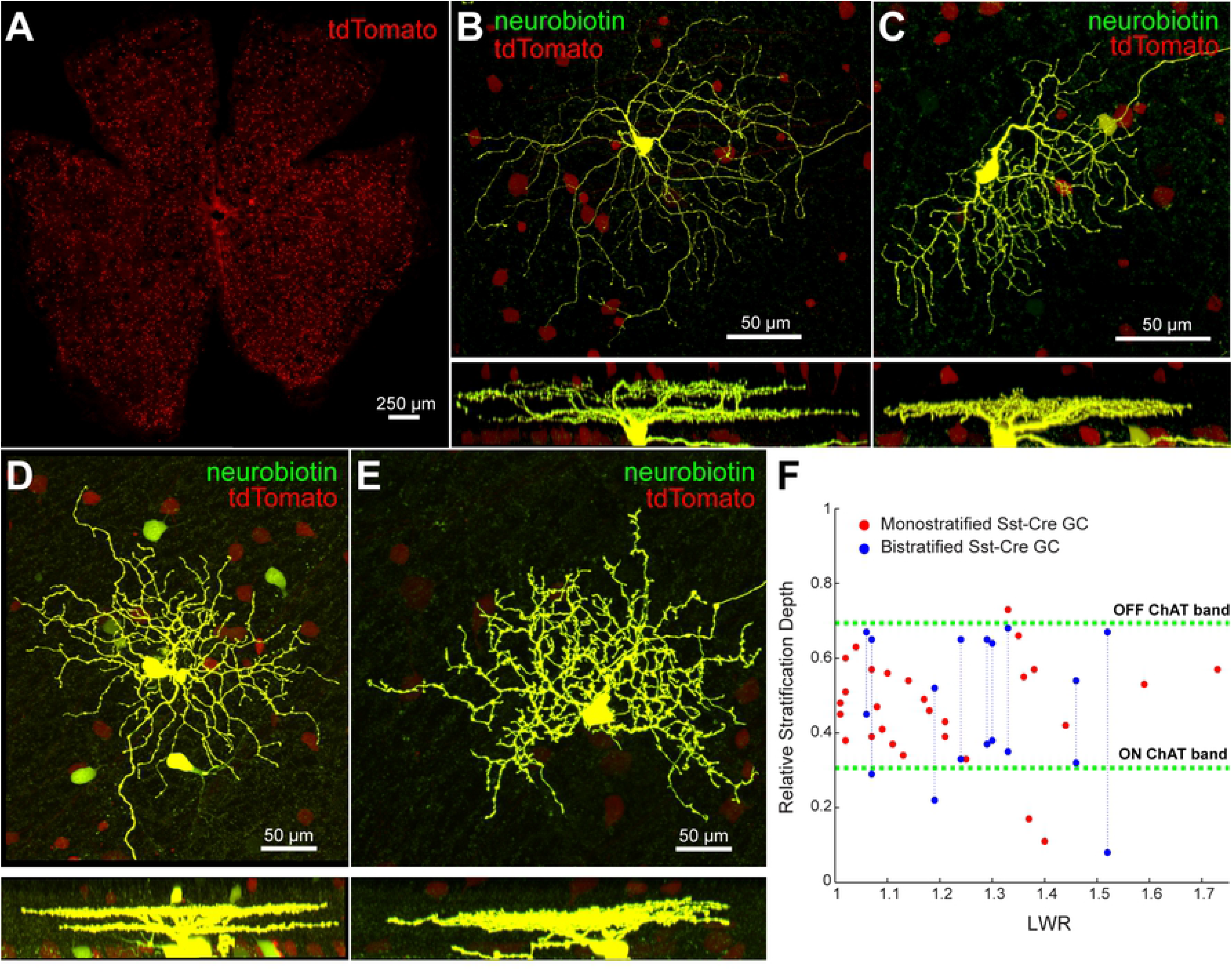
Morphological analysis of SST-*ires*-Cre positive GCs shows a diverse subset of cells. A. Confocal fluorescence image of a wholemount SST-*ires*-Cre x Ai9 mouse retina. SST-*ires*-Cre positive neurons express tdTomato when crossed with an Ai9 reporter mouse. Fluorescence expression was dependent on the genotype of the mouse, with homozygous animals expressing tdTomato in 50% more cells in the GC layer compared with heterozygous animals. B. Example of an SST-*ires*-Cre positive bistratified GC, filled with neurobiotin using a sharp electrode and cross-reacted with streptavidin-conjugated Alexa 488. Side view (below) shows dendrites passing between the ON and OFF layers in the IPL, indicating this may be the previously reported SBC GC. C. Example of an SST-*ires*-Cre positive monostratified GC, visualized as in B. Strong dendritic asymmetry predicts for this GC an orientation-selective (OS) visual response. D. Example of an SST-*ires*-Cre positive bistratified GC with 5-6 neurobiotin-positive cell bodies near the primary (injected) GC. Apparent gap-junction coupled cells surrounding the injected GC were commonly observed following SST-*ires*-Cre positive GC fills. Absence of an axon in the surrounding labeled cells indicates that the secondary-labeled cells were all amacrine cells. E. Example of an SST-*ires*-Cre positive monostratified GC with large dendritic arbor (compare with cell in panel C). F. Scatter plot of length-to-width ratio (LWR) vs. stratification depth of SST-*ires*-Cre positive GCs visualized with a confocal microscope following targeted neurobiotin fills with sharp electrodes, or with two-photon fluorescence imaging following targeted whole-cell recording.

Whether homozygous animals labeled specific GC types that were not labeled in heterozygous animals, or whether the increase in the number of labeled cells simply reflected an increase in the labeled fraction of a fixed set of GC types remained unclear and data obtained from both genotypes were combined in this study.

Morphological reconstruction following neurobiotin fills of genetically labeled GCs in Sst-Cre x Ai9 mice showed that Sst-Cre GCs are a diverse but limited subset of GC types (Figure 1B-E). For example, labeled types never included the large soma, alpha-type GCs, but did include bistratified GCs and monostratified GCs with dendritic arbors terminating in the ON and OFF layers of the inner plexiform layer (IPL; Figure 1F). In some instances, neurobiotin fills resulted in additional labeling of nearby cell bodies. These cells, all apparent ACs, were labeled through gap junction coupling with the neurobiotin-filled Sst-Cre GC (Fig 1D). Quantitative analysis of stratification depth showed that many Sst-Cre positive GCs stratified between the ChAT bands, indicating that the labeled population was non-direction selective and received predominant synaptic input from bipolar cell types with axon terminals in the central IPL, only (Figure 1F). Quantitative analysis of length-to-width ratio (L:W) showed that the dendritic arbors of a subset of Sst-Cre positive GCs was distinctly asymmetrical (L:W ratio < 1.33; Figure 1F, example GC shown in Figure 1C). This asymmetric dendritic morphology predicts that at least some Sst-Cre positive GCs are intrinsically orientation selective, which we confirm below.

### Calcium Imaging Reveals DS, OS and SBC Sst-Cre Positive GCs

Next, we used two-photon fluorescence calcium imaging to study visually evoked responses of Sst-Cre positive GCs at the neuron population level. To obtain GCaMP6f expression in Sst-Cre GCs, we crossed Sst-Cre mice with the Ai95 reporter mouse (Madisen et al., 2015). The resulting fluorescence expression pattern matched that of the Sst-Cre x Ai9 cross used in the anatomical analyses (above). We recorded GCaMP6f fluorescence responses of local GC populations comprising 2-10 labeled cells (80 × 80 μm field of view) to a series of photopic visual stimuli (1.2 · 10^5^ R^∗^ / rod and cone/s) projected onto the photoreceptor layer of a whole-mount retina preparation. We used the modulation amplitude of each cell’s response to oriented drifting sinewave gratings (Fig. 2A-C) to classify labeled GCs into three broad classes: direction selective (DS; DS index, DSi ≥ 0.4), orientation selective (OS; OS index, OSi ≥ 0.4 and DSi < 0.4), and neither DS nor OS (non-DS/non-OS; OSi < 0.4 and DSi < 0.4).

**Fig 2.**
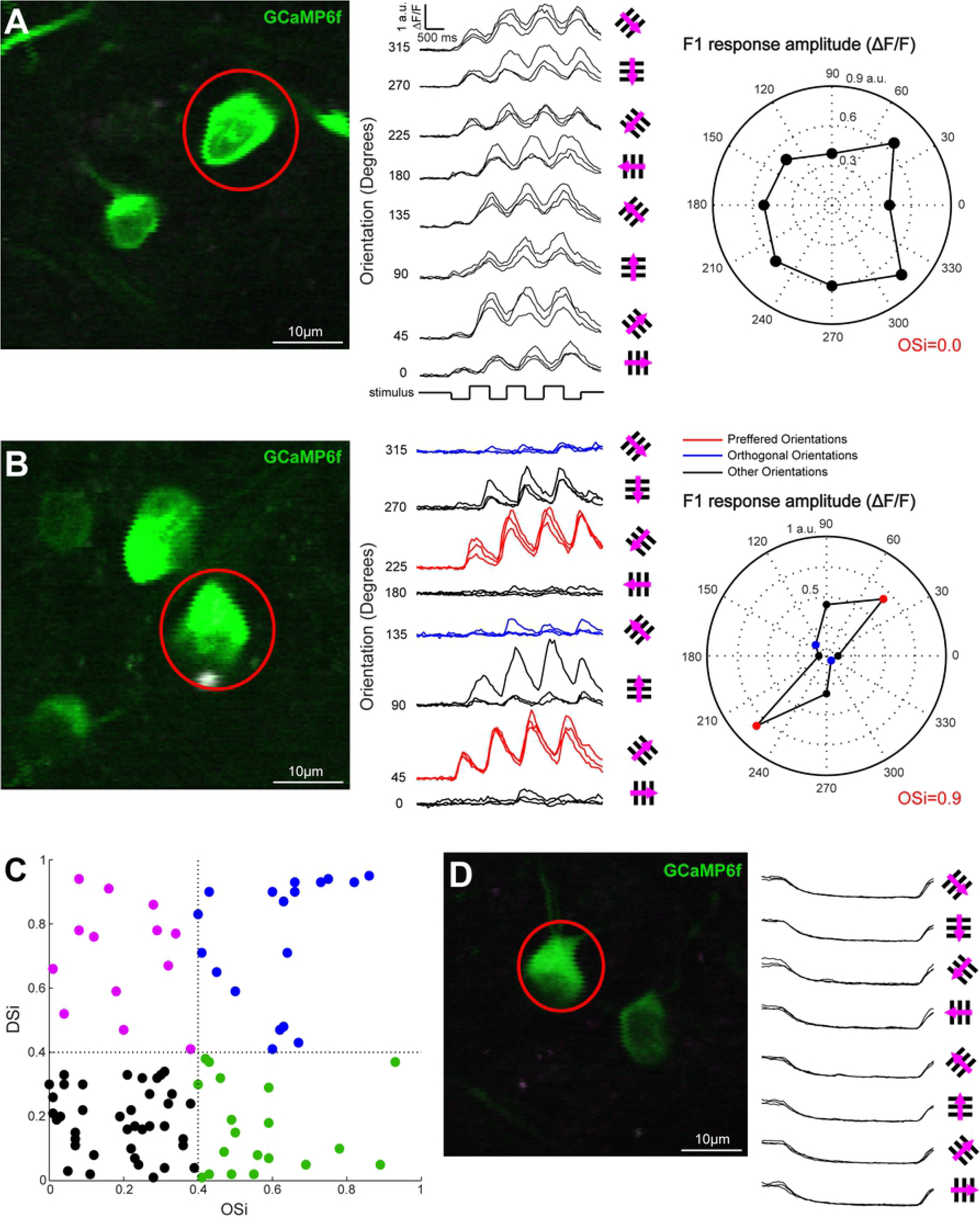
GCaMP6f calcium imaging of SST-*ires*-Cre x Ai95 GCs identifies DS, OS and SBC GCs. A. Example non-DS, non-OS GC with strong responses to 8 different orientations of drifting gratings. Calcium responses to the drifting gratings (middle) are shown for the cell body circled in red in the two-photon image (left). Fluorescent responses were analyzed using a Fast-fourrier transform (FFT); polar plot (right) shows the F1 response to each of the 8 directions. OSi of this example GC is 0.0. B. Example OS GC with a strong direction preference, OSi = 0.9. Preferred axis responses are shown in red while the orthogonal axis is shown in blue. C. DSi versus OSi plot of GCaMP6f responses in SST-*ires*-Cre positive GCs. Overall population responses indicate a heterogeneous set of GCs that include non-DS/non-OS GCs (black dots), DS GCs (magenta and blue dots) and OS GCs (green dots). D. Example GC with strong suppression of spontaneous activity following presentation of drifting gratings of any orientation. This RGC apparently corresponds to the SBC GC previously described (see Introduction).

Of the recorded cells, the largest group were non-DS/non-OS (44.3%; n = 39 / 88 total cells; Fig. 2D, black dots). About one third of the recorded cells classified as DS (34%; n = 30 / 88; Fig. 2D, magenta and blue dots). The remaining recorded cells classified as OS (21.6%; n = 19 / 88; Fig. 2D, green dots).

Calcium imaging at the population level consistently revealed a GC type whose calcium response was strongly suppressed during all directions of motion of the drifting grating stimulus. These cells showed substantial fluorescence during presentation of a mid-level grey background prior to grating onset, suggesting tonic excitation to a static and spatially uniform background. High baseline fluorescence in these cells was reduced following the onset of all eight orientations of our drifting grating stimulus, suggesting tonic suppression of the cells’ responses during the stimulus presentation (Fig 2E). This specific visually-evoked response pattern matches that of the previously described suppressed by contrast GC (Tien et al., 2015). Moreover, our set of GC morphologies obtained with neurobiotin fills includes a type with dendrites that appear to pass between the ON and OFF layers of the IPL (Figure 1B), similar to the reported morphology of the SBC cell (Tien et al., 2015).

### Electrophysiological recording shows four primary Sst-Cre positive GC types

To assess the response properties of Sst-Cre positive GCs in further detail, we targeted individual cells for whole-cell recording during visual stimulation of the retina with contrast reversing spots, and drifting and static gratings. We measured for each cell the current-voltage relationship (IV curves), conductance, and DSi and OSi. We distinguished four GC response types based on the shape and relative magnitude of their excitatory and inhibitory conductance, recorded in voltage clamp at the reversal potential for chloride and cations, respectively (V_hold_: E_Cl_ = −69 mV, E_cat_ = 15 mV; Fig. 3).

**Fig 3.**
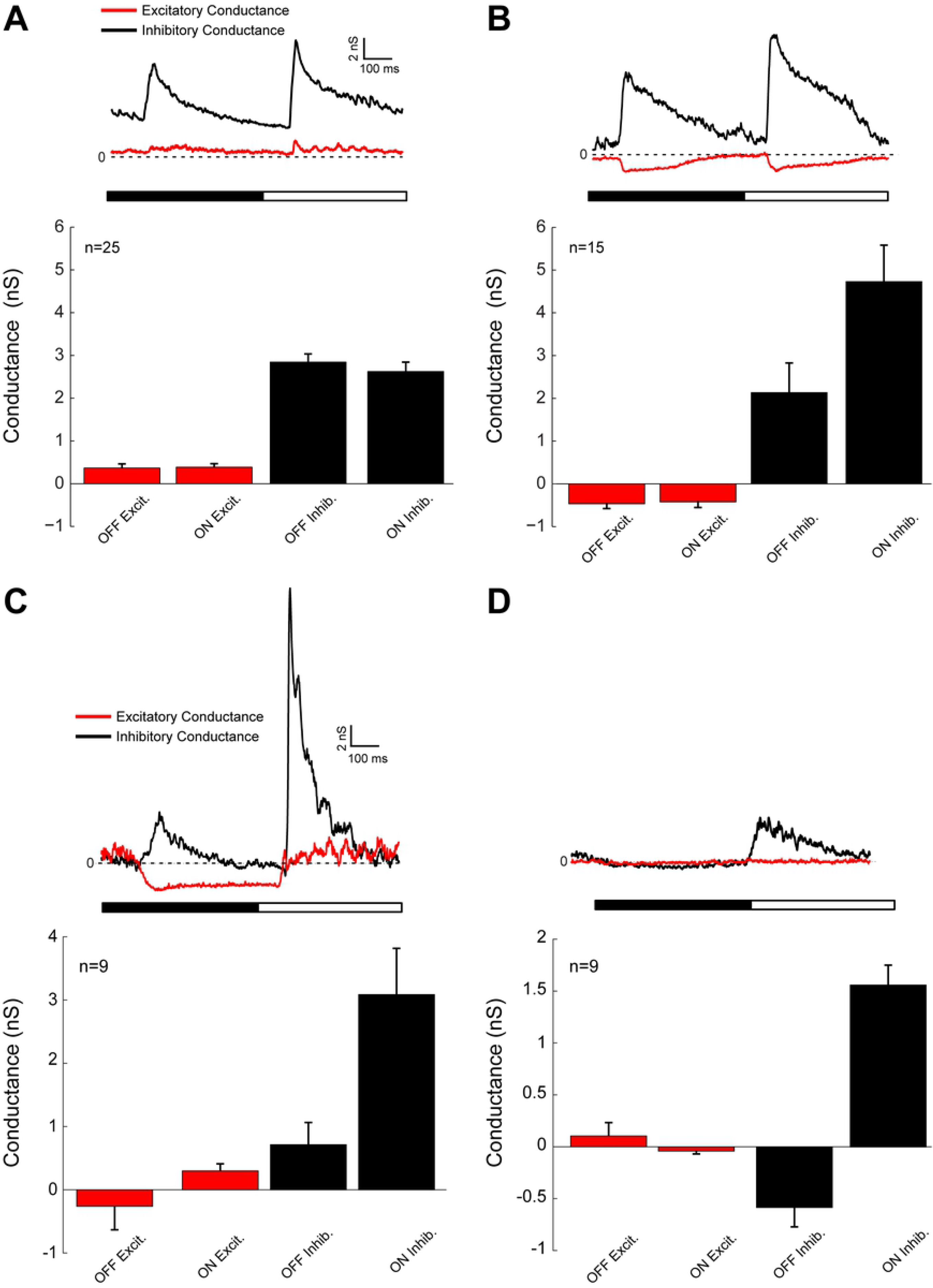
Whole-cell recording reveals four commonly recorded subtypes of SST-*ires*-Cre positive GCs. A. Example conductance (top) and population average conductance (bottom) for the most commonly recorded Sst-Cre positive GC type (n=25/149 GCs total). Inhibitory conductances are shown in black and excitatory conductances are shown in red for all traces and graphs. B. Example conductance (top) and average conductances (bottom) for the second most commonly recorded RGC (n=15/149). This RGC subtype shows strong OS and bistratified morphology, similar to a previously reported OS GCs (see Introduction). C. Example conductance (top) and average conductances (bottom) for an apparent SBC GC. This GC showed strong suppression of excitatory conductance during light decrements and strong inhibitory conductance during both light increments and decrements. D. Example conductance (top) and average conductances (bottom) of the fourth commonly recorded RGC in the SST-*ires*-Cre mouse. The most prominent feature of this GC is the strong inhibitory conductance following a light increment and little to no change in excitatory conductance following either a light increment or decrement.

Most Sst-Cre positive cells showed a large inhibitory conductance, typically at both onset and offset, and a relatively small excitatory conductance (Fig. 3). Example conductances and the average ON and OFF excitatory and inhibitory conductances are shown for the four most frequently recorded Sst-Cre GC response types (Figure 3A-D). These GC types accounted for 38.9% of the total number of GCs recorded (n=58/149), comprising 16.7% (Figure 3A), 10% (Figure 3B), 6% (Figure 3C), and 6% (Figure 3D) of the recorded population.

The electrophysiological recordings identified a cell type that was distinctly suppressed by contrast, consistent with the calcium imaging experiments. Similar to the SBC in a previous report (Tien et al., 2015), this cell type was characterized by an increase in inhibitory conductance to light decrements and massive inhibition during light increments, along with suppression of excitatory currents during light decrements (Figure 3C).

Analysis of membrane voltage and excitatory and inhibitory currents in response to drifting gratings further revealed a GC type that was strongly orientation-tuned (Figure 4A, B; B, example OSi = 0.76). Both the excitatory and inhibitory currents reflected the OS of the membrane voltage response, but OS of the excitatory input was typically dominant. We found a similar trend when comparing the OSi of the excitatory currents to the OSi of the inhibitory currents (Figure 4C). When plotting excitatory current OSi versus inhibitory current OSi, the majority of the data points lie to the left/above the unity line, i.e., where data points would fall if OSi values of both currents were equal (Figure 4C). Instead, a linear fit to the data shows a slope of 1.26, indicating greater OSi of the excitatory current compared with OSi of the inhibitory current recorded in the same cell. This suggests that the excitatory input to GCs may drive OS to a larger degree than inhibitory currents. This is intriguing, and could reflect oriented excitatory input, for example from elongated dendritic morphology.

**Fig 4.**
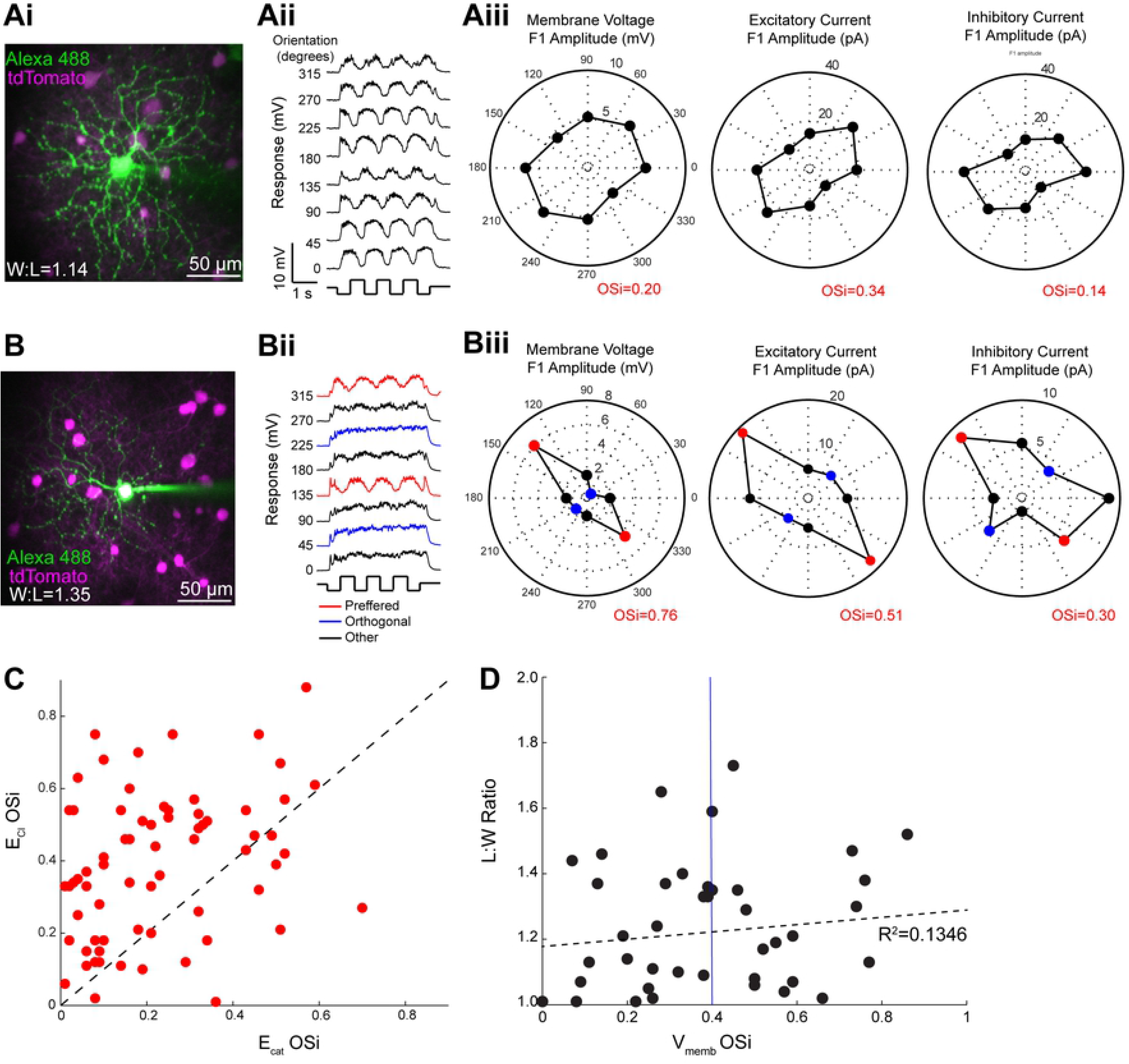
Excitation is the primary driver of OS responses. A. Z-projection of a two-photon fluorescence image stack of a non-OS GC, die-filled during whole-cell recording (Ai). Average membrane voltage responses (Aii) to drifting gratings at 8 orientations. Polar plots (Aiii) of membrane voltage, excitatory current, and inhibitory current (left through right) for the cell shown in Ai. B. Z-projection of a two-photon fluorescence image stack of an OS GC (Bi), die-filled during whole-cell recording. Bii and Biii, same as in A. C. Comparison of the OSi computed from inhibitory currents (E_cat,_ x-axis) and excitatory currents (E_Cl_, y-axis) of SST-*ires*-Cre positive GCs obtained with targeted whole-cell recording. In the majority of cells, OSi of the excitatory current exceeded that of the inhibitory current. Unity line, dashed black. D. Comparison of the OSi (x-axis) based on the membrane voltage response vs. the L:W ratio (y-axis) of SST-*ires*-Cre positive GCs. The black dashed line represents the line of best fit and shows poor correlation between the OSi and the L:W ratio (R^2^=0.1346), indicating that OS does not result from asymmetric morphology.

To test this, we compared the L:W ratio (length of the dendritic arbor at its greatest spatial extent, divided by the length of the orthogonal extent) of recorded GCs to their respective OSi values. We found a weak correlation at best between L:W ratio and OSi (Figure 4D). Indeed, linear regression analysis showed that just 13.46% of the OSi variation can be accounted for by the size of the L:W ratio. This suggests that OS in these GCs is an emergent, circuit-level property rather than a consequence of asymmetric dendritic morphology. The neural underpinnings of this emergent OS are beyond the scope of this work, but may now be efficiently addressed through targeted recording in future studies.

## Discussion

We characterized fluorescently labeled GCs in the Sst-Cre mouse crossed with transgenic fluorescent reporter lines. We report that the labeled cell population is relatively sparse, comprising just 4-6% of all GCs in the mouse retina. The number of labeled cells is increased in homozygous compared with heterozygous Sst-Cre animals, indicating low-level Cre expression in potentially many retinal cells in this line whereas substantial Cre-expression in the ganglion cell layer is limited to a specific subset of GC types. This makes the Sst-Cre transgenic mouse line useful for studies of identified GC circuits within in the mouse retina.

Using calcium imaging, we distinguished among the labeled GC population direction selective, orientation selective, and suppressed by contrast-type GCs. Targeted electrophysiological whole-cell recording of excitatory and inhibitory currents showed DS, OS and suppressed-by-contrast type responses consistent with the morphological and calcium imaging data, with a dominant role for inhibition in each type. OS and SBC GCs account for two of the four Sst-Cre GCs most commonly recorded in our experiments. We currently lack a detailed understanding of the presynaptic circuit and cell-intrinsic mechanisms that generate the distinct visual responses of these cell types. We anticipate that the Sst-Cre transgenic mouse line will be instrumental for studying the SBC and the OS GC that has been previously characterized exclusively through whole-cell recording of unlabeled GCs (Nath and Schwartz, 2017).

The utility of cell type-selective Cre-recombinase gene expression is not easily overstated. For example, the ability to express fluorescent sensor proteins by crossing with the Ai95 (GCaMP6f) reporter mouse allows for efficient studies and analyses at the population level of one or a small number of identified GC types. Combined with real-time analysis of fluorescence responses obtained during visual stimulation, targeted recording of cells based on these calcium responses strongly increases the efficiency with which the OS or SBC GC may be studied electrophysiologically and morphologically. We conclude that the Sst-Cre mouse represents a currently underutilized tool for circuit-level studies of the mouse retina that is poised to provide new insights into a diverse subset of functionally identified GCs.

## Acknowledgements

This research was funded by NIH/NEI grant EY028188 (BGB) and the E. Matilda Ziegler Foundation (BGB).

## References

Baden T, Berens P, Franke K, Roman Roson M, Bethge M, Euler T (2016) The functional diversity of retinal ganglion cells in the mouse. Nature 529:345–350.

Buffelli M, Burgess RW, Feng G, Lobe CG, Lichtman JW, Sanes JR (2003) Genetic evidence that relative synaptic efficacy biases the outcome of synaptic competition. Nature 424:430–434.

Fransen JW, Borghuis BG (2017) Temporally Diverse Excitation Generates Direction-Selective Responses in ON- and OFF-Type Retinal Starburst Amacrine Cells. Cell Rep 18:1356–1365.

Huberman AD, Wei W, Elstrott J, Stafford BK, Feller MB, Barres BA (2009) Genetic identification of an On-Off direction-selective retinal ganglion cell subtype reveals a layer-specific subcortical map of posterior motion. Neuron 62:327–334.

Larsen JN, Bersani M, Olcese J, Holst JJ, Moller M (1990) Somatostatin and prosomatostatin in the retina of the rat: an immunohistochemical, in-situ hybridization, and chromatographic study. Vis Neurosci 5:441–452.

Madisen L, Zwingman TA, Sunkin SM, Oh SW, Zariwala HA, Gu H, Ng LL, Palmiter RD, Hawrylycz MJ, Jones AR, Lein ES, Zeng H (2010) A robust and high-throughput Cre reporting and characterization system for the whole mouse brain. Nat Neurosci 13:133–140.

Madisen L et al. (2015) Transgenic mice for intersectional targeting of neural sensors and effectors with high specificity and performance. Neuron 85:942–958.

Martersteck EM, Hirokawa KE, Evarts M, Bernard A, Duan X, Li Y, Ng L, Oh SW, Ouellette B, Royall JJ, Stoecklin M, Wang Q, Zeng H, Sanes JR, Harris JA (2017) Diverse Central Projection Patterns of Retinal Ganglion Cells. Cell Rep 18:2058–2072.

Nath A, Schwartz GW (2017) Electrical synapses convey orientation selectivity in the mouse retina. Nat Commun 8:2025.

Park SJ, Kim IJ, Looger LL, Demb JB, Borghuis BG (2014) Excitatory synaptic inputs to mouse on-off direction-selective retinal ganglion cells lack direction tuning. J Neurosci 34:3976–3981.

Park SJ, Borghuis BG, Rahmani P, Zeng Q, Kim IJ, Demb JB (2015) Function and Circuitry of VIP+ Interneurons in the Mouse Retina. J Neurosci 35:10685–10700.

Pologruto TA, Sabatini BL, Svoboda K (2003) ScanImage: flexible software for operating laser scanning microscopes. Biomed Eng Online 2:13.

Rickman DW, Blanks JC, Brecha NC (1996) Somatostatin-immunoreactive neurons in the adult rabbit retina. J Comp Neurol 365:491–503.

Sanes JR, Masland RH (2015) The types of retinal ganglion cells: current status and implications for neuronal classification. Annu Rev Neurosci 38:221–246.

Scheyltjens I, Arckens L (2016) The Current Status of Somatostatin-Interneurons in Inhibitory Control of Brain Function and Plasticity. Neural Plast 2016:8723623.

Sivyer B, Taylor WR, Vaney DI (2010) Uniformity detector retinal ganglion cells fire complex spikes and receive only light-evoked inhibition. Proc Natl Acad Sci U S A 107:5628–5633.

Sumbul U, Song S, McCulloch K, Becker M, Lin B, Sanes JR, Masland RH, Seung HS (2014) A genetic and computational approach to structurally classify neuronal types. Nat Commun 5:3512.

Taniguchi H, He M, Wu P, Kim S, Paik R, Sugino K, Kvitsiani D, Fu Y, Lu J, Lin Y, Miyoshi G, Shima Y, Fishell G, Nelson SB, Huang ZJ (2011) A resource of Cre driver lines for genetic targeting of GABAergic neurons in cerebral cortex. Neuron 71:995–1013.

Tien NW, Pearson JT, Heller CR, Demas J, Kerschensteiner D (2015) Genetically Identified Suppressed-by-Contrast Retinal Ganglion Cells Reliably Signal Self-Generated Visual Stimuli. J Neurosci 35:10815–10820.

White CA, Chalupa LM, Johnson D, Brecha NC (1990) Somatostatin-immunoreactive cells in the adult cat retina. J Comp Neurol 293:134–150.

Williams RW, Strom RC, Rice DS, Goldowitz D (1996) Genetic and environmental control of variation in retinal ganglion cell number in mice. J Neurosci 16:7193–7205.

Yonehara K, Balint K, Noda M, Nagel G, Bamberg E, Roska B (2011) Spatially asymmetric reorganization of inhibition establishes a motion-sensitive circuit. Nature 469:407–410.

